# Fitness barriers limit reversion of a proofreading-deficient coronavirus

**DOI:** 10.1101/618249

**Authors:** Kevin W. Graepel, Maria L. Agostini, Xiaotao Lu, Nicole R. Sexton, Mark R. Denison

## Abstract

The 3′-to-5′ exoribonuclease in coronavirus (CoV) nonstructural protein 14 (nsp14-ExoN) mediates RNA proofreading during genome replication. ExoN catalytic residues are arranged in three motifs: I (DE), II (E), III (D). Alanine substitution of the motif I residues (AA-E-D, four nucleotide substitutions) in murine hepatitis virus (MHV) and SARS-CoV yields viable mutants with impaired replication and fitness, increased mutation rates, and attenuated virulence *in vivo*. Despite these impairments, MHV- and SARS-CoV ExoN motif I AA mutants (ExoN-AA) have not reverted at motif I in diverse *in vitro* and *in vivo* environments, suggesting that profound fitness barriers prevent motif I reversion. To test this hypothesis, we engineered MHV-ExoN-AA with 1, 2 or 3 nucleotide mutations along genetic pathways to AA-to-DE reversion. We show that engineered intermediate revertants were viable but had no increased replication or competitive fitness compared to MHV-ExoN-AA. In contrast, a low passage (P10) MHV-ExoN-AA showed increased replication and competitive fitness without reversion of ExoN-AA. Finally, engineered reversion of ExoN-AA to ExoN-DE in the presence of ExoN-AA passage-adaptive mutations resulted in significant fitness loss. These results demonstrate that while reversion is possible, at least one alternative adaptive pathway is more rapidly advantageous than intermediate revertants and may alter the genetic background to render reversion detrimental to fitness. Our results provide an evolutionary rationale for lack of ExoN-AA reversion, illuminate potential multi-protein replicase interactions and coevolution, and support future studies aimed at stabilizing attenuated CoV ExoN-AA mutants.

**IMPORTANCE:** Coronaviruses encode an exoribonuclease (ExoN) that is important for viral replication, fitness, and virulence, yet coronaviruses with a defective ExoN (ExoN-AA) have not reverted under diverse experimental conditions. In this study, we identify multiple impediments to MHV-ExoN-AA reversion. We show that ExoN-AA reversion is possible but evolutionarily unfavorable. Instead, compensatory mutations outside of ExoN-AA motif I are more accessible and beneficial than partial reversion. We also show that coevolution between replicase proteins over long-term passage partially compensates for ExoN-AA motif I but renders the virus inhospitable to a reverted ExoN. Our results reveal the evolutionary basis for the genetic stability of ExoN-inactivating mutations, illuminate complex functional and evolutionary relationships between coronavirus replicase proteins, and identify potential mechanisms for stabilization of ExoN-AA coronavirus mutants.

## INTRODUCTION

The rapid evolution of RNA viruses represents a significant challenge for preventing, treating, and eradicating RNA viral diseases. High mutation rates in RNA viruses generate extensive opportunities to overcome evolutionary hurdles, such as antiviral drugs, host immunity, or engineered attenuating changes (1). The evolutionary pathways traversed by RNA viruses are shaped by natural selection, which will favor some evolutionary trajectories more than others based on whether mutations are beneficial, deleterious, or neutral (2). Predicting the likely results of RNA virus evolution is an important step for anticipating viral emergence and for developing escape-resistant antiviral drugs and vaccines (3, 4).

Coronaviruses (CoVs) are a family of positive-sense RNA viruses that cause human illnesses ranging from the common cold to severe and lethal respiratory disease (5). All CoVs encode a proofreading exoribonuclease within nonstructural protein 14 (nsp14-ExoN) that is critical for replication, fidelity, fitness, and virulence, and ExoN-inactivation has been proposed as a strategy for live-attenuated vaccine development (6–15). As members of the DEDDh superfamily of exonucleases, CoV ExoNs hydrolyze nucleotides using four metal-coordinating amino acids arranged in 3 motifs: I (DE), II (E), III (D) (16, 17). Alanine substitution of ExoN motif I (DE-to-AA) disrupts ExoN biochemical activity in both SARS-CoV and human CoV 229E (hCoV-229E) (16, 18, 19). The betacoronaviruses murine hepatitis virus (MHV) and SARS-CoV tolerate disruption of ExoN activity [ExoN(-)] but display mutator phenotypes accompanied by defects in replication, competitive fitness, and evasion of innate immune responses (10, 13, 14). ExoN active site mutants in alphacoronaviruses, including transmissible gastroenteritis virus and hCoV-229E, have yet to be recovered and are proposed to be lethal for replication (19, 20).

Given the critical role of ExoN in CoV biology and the elevated mutation rate, we expected that natural selection would repeatedly drive reversion of the ExoN-inactivating substitutions. In line with this expectation, ExoN motif III mutants of SARS-CoV and MHV rapidly and repeatedly revert ((14) and unpublished observations). In contrast, we have never detected partial or complete reversion of ExoN motif I mutants (ExoN-AA) in SARS-CoV or MHV during 10 years of study and hundreds of experiments. More specifically, we have not detected consensus or minority variants of any kind at the motif I AA codons in either virus strain during acute infections and prolonged passage in tissue culture and following treatment with multiple nucleoside analogues (6–11, 13, 14). SARS-CoV-ExoN-AA also is stable during acute and persistent animal infections in immunocompetent and immune-compromised mice (12). The lack of primary reversion is not due simply to reduced adaptive capacity, as both SARS-CoV- and MHV-ExoN-AA can adapt for increased replication (7, 14). Most strikingly, long-term passage of MHV-ExoN-AA (250 passages, P250) yielded a highly fit population that had directly compensated for defective proofreading through evolution of a likely high-fidelity RdRp (7). Yet, where primary reversion would have required just four total consensus mutations, MHV-ExoN-AA-P250 contained more than 170.

In this study, we sought to determine whether specific genetic or fitness barriers prevent primary reversion of ExoN motif I AA. To this end, we identified and engineered viable genetic pathways towards ExoN-AA motif I reversion in MHV (hereafter, ExoN-AA). Our results show that partial reversion did not confer a selective advantage compared to ExoN-AA. Further, ExoN-AA adapted within 10 passages for greater fitness than any of the intermediate revertants. Finally, restoration of WT-ExoN-DE in the setting of passage-selected mutations in the nsp12 RNA-dependent RNA polymerase (RdRp) and nsp14-ExoN exacted profound fitness costs. Together, these data are the first observation of an ExoN(-) CoV genotype-fitness landscape and identify multiple genetic barriers underlying ExoN(-) motif I stability in MHV. Further, they suggest extensive coevolution between MHV replicase proteins during adaptation and reveal potential strategies for stabilizing ExoN mutant CoVs.

## RESULTS

### Primary reversion of ExoN(-) motif I

MHV-ExoN(-), hereafter ExoN-AA, contains two engineered substitutions in each codon of motif I, such that complete reversion to WT-ExoN-DE requires mutations to all four sites (Figure 1A). Viral mutation rates in the absence of proofreading range from 10^−4^ to 10^−6^ mutations per nucleotide per round of replication (μ) (1). Assuming an ExoN-AA mutation rate of 10^−4^ μ and accounting for codon degeneracy, the probability of restoring the native amino acid sequence in a single round of replication is 10^−18^. Only rarely do ExoN-AA titers exceed 10^6^ PFU/mL, so it is unlikely that ExoN-AA could navigate this genetic barrier in a single infectious cycle. Thus, we hypothesized that ExoN-AA reversion, if possible, would proceed incrementally. To identify potential pathways towards ExoN-AA reversion, we examined the possible single-nucleotide substitutions surrounding A89 and A91 (Figure 1B). Three mutations are synonymous, and five mutations yield amino acids unlikely to coordinate with the positively-charged metals required for ExoN catalysis (glycine, valine, proline, threonine, and serine) (16, 19, 21, 22). One mutation per site can restore the acidic charge (i.e. AA-to-ED) but not the native amino acid. These variants have not been tested in a CoV ExoN, but biochemical studies of *E. coli* DNA polymerase I ExoN mutants suggest that these conservative substitutions would not restore WT-like ExoN activity (23). We predicted stepwise pathways to ExoN-AA→DE reversion based on restoration of acidic charge followed by reversion to native amino acids (Figure 1C). We engineered and recovered variants in ExoN-AA requiring three mutations (3nt; ExoN-AD, ExoN-EA), two mutations (2nt; ExoN-DA, ExoN-ED, ExoN-AE), or one mutation (1nt; ExoN-DD, ExoN-EE) for reversion to WT-ExoN-DE (Table 1). We will hereafter refer to these mutants as intermediate revertants. All intermediate revertants generated viable progeny during recovery, demonstrating that reversion to WT-ExoN-DE along these pathways is theoretically possible. The 3nt and 2nt mutants were genetically stable during recovery, as confirmed by dideoxy sequencing. However, both 1nt mutants (ExoN-DD and ExoN-EE) reverted to WT-ExoN-DE during three independent recovery attempts, suggesting that these two variants are less fit than WT-ExoN-DE and demonstrating that reversion by 1nt mutation is readily accessible. To test whether the 3nt or 2nt mutants would revert more rapidly than ExoN-AA (4nt), we passaged three lineages of each mutant 10 times at multiplicities of infection (MOI) of 0.5 and 0.01 PFU/cell. We harvested supernatants and screened for reversion by visual inspection of plaque phenotypes at each passage. WT-ExoN-DE and WT-like viruses produce uniform, large plaques, while ExoN-AA-like viruses yield small, variably-sized plaques (13). When we observed mixed plaque phenotypes, we sequenced three large plaques from each lineage to confirm reversion. The 3nt (ExoN-AD and ExoN-EA) and 2nt (ExoN-DA and ExoN-ED) intermediate revertants showed no evidence of reversion over 10 passages at either MOI (Table 1). In contrast, the 2nt ExoN-AE contained WT-revertants by P2 in all lineages at MOI = 0.5 PFU/cell and by P8 in one lineage at MOI = 0.01 PFU/cell. Once observed, WT-revertants dominated the ExoN-AE population for the remaining passages. These data indicate that at least one 2nt mutation pathway can lead to full reversion in tissue culture. The probability of ExoN-AE arising during a single infectious cycle of ExoN-AA is low but theoretically achievable (~10^−9^), so ExoN-AA could conceivably revert within just two infectious cycles. However, complete reversion has never been observed even during prolonged passage or persistent infections, suggesting that additional barriers to the replication, fitness, or maintenance of intermediate revertants exist.

**Table 1.** Recovery and passage of intermediate revertants.

| Virus | # of mutations to<br>WT-ExoN-DE | Motif I sequence<sup>a</sup> | # of reverted lineages by passage 10<sup>b</sup> |  |
| --- | --- | --- | --- | --- |
|  |  |  | MOI = 0.01 | MOI = 0.5 |
| ExoN-AA | 4 | G**CA**...G**CT** | 0/3 | 0/3 |
| ExoN-AD | 3 | G**CA**...G**AT** | 0/3 | 0/3 |
| ExoN-EA | 3 | G**AA**...G**CT** | 0/3 | 0/3 |
| ExoN-DA | 2 | GAT...G**CT** | 0/3 | 0/3 |
| ExoN-AE | 2 | G**CA**...GAA | 1/3 (by P8) | 3/3 (by P2) |
| ExoN-ED | 2 | G**AA**...G**AT** | 0/3 | 0/3 |
| ExoN-EE | 1 | G**AA**...GAA | n.d. | n.d. |
| ExoN-DD | 1 | GAT...G**AT** | n.d. | n.d. |
| WT-ExoN-DE | 0 | GAT...GAA | n.d. | n.d. |

**Figure 1.**
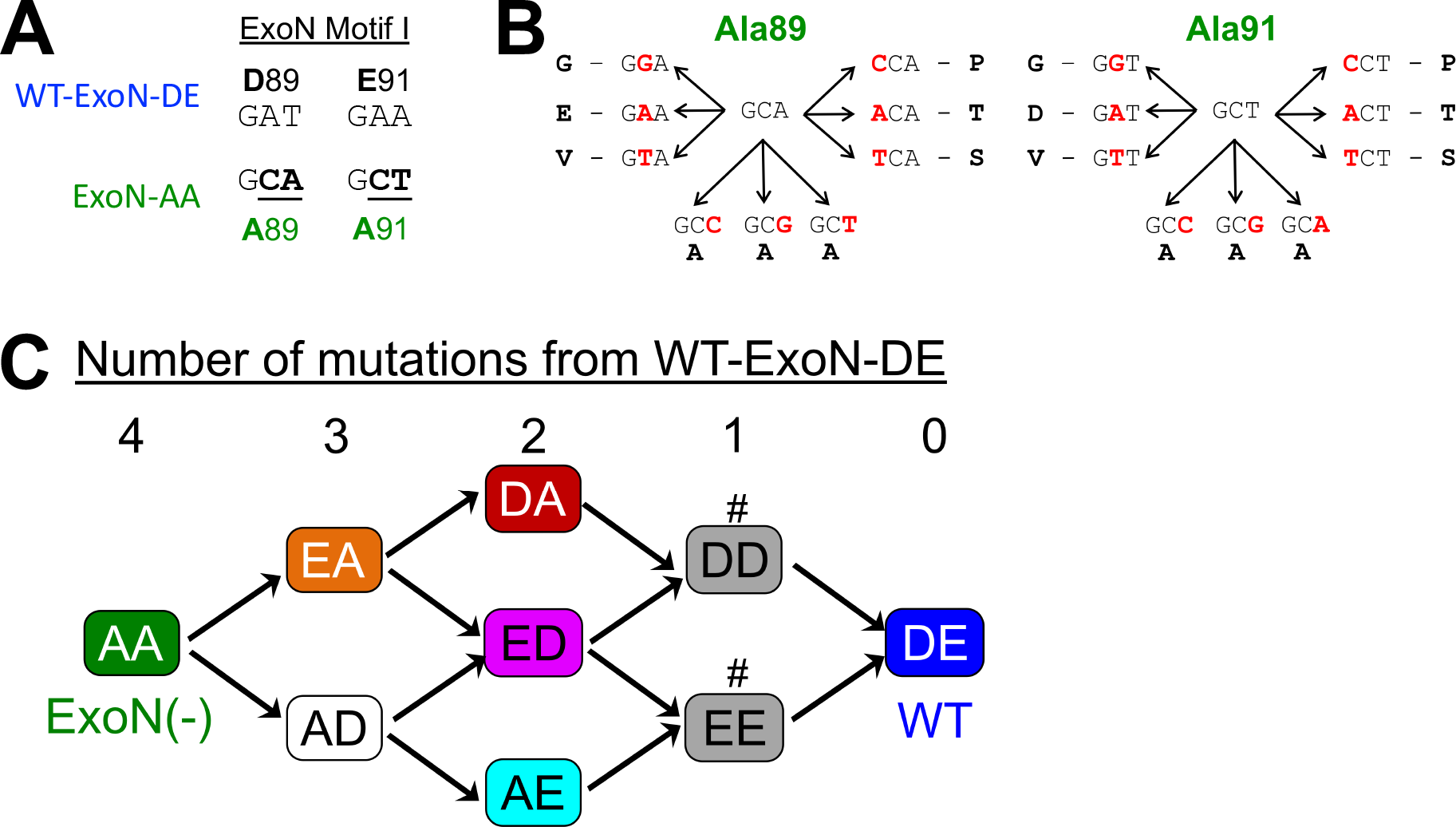
Sequence landscape around ExoN-AA motif I. (*A*) ExoN motif I nucleotide sequences. (*B*) Landscape of single-nucleotide substitutions within ExoN-AA motif I. (*C*) Predicted pathways to reversion of ExoN-AA. Variants marked with # reverted to WT during three independent recovery attempts.

### Partial reversion of MHV-ExoN(-) motif I does not confer a selective advantage

Because the intermediate revertants are viable as recombinants but are not found in ExoN-AA populations, we hypothesized that they confer no selective advantage over ExoN-AA (8, 9, 13). To test this hypothesis, we first analyzed replication of the 3nt and 2nt intermediate revertants (Figure 2A). All variants achieved similar peak titers to ExoN-AA, but detailed examination of their kinetics suggested a potential delay of up to 1.5 hours for all intermediate revertants compared to ExoN-AA. Of note, ExoN-AE was the most delayed, and we detected WT-ExoN-DE revertants in two of three replicates, suggesting increased selective pressure against this variant. We next measured the competitive fitness of each intermediate revertant relative to a recombinant ExoN-AA containing seven silent mutations in the nsp2 coding region (ExoN-AA-reference). Intermediate revertants were mixed with an equal titer of ExoN-AA-reference at a combined MOI = 0.05 PFU/cell and passaged four times. The ratio of each intermediate revertant to ExoN-AA-reference was quantified at each passage by RT-qPCR, and the change in ratio over time was used to calculate their relative fitness. WT-ExoN-DE was significantly more fit than ExoN-AA, whereas the intermediate revertants (ExoN-AD, -EA, -DA, and -ED) had no increased fitness relative to ExoN-AA (Figure 2B). The apparent increased fitness of ExoN-AE resulted from all lineages reverting to WT-ExoN-DE during the experiment. Finally, our previous studies have shown that adaptation of ExoN-AA includes partial compensation for the replication fidelity defect, as measured by reduced susceptibility to the mutagen 5-fluorouracil (5-FU) (7–11, 24). None of the intermediate variants demonstrated statistically significant differences in 5-FU sensitivity as compared to ExoN-AA (Figure 2C). Thus, with the exception of the ExoN-AE→DE revertants, no 3nt and 2nt intermediate genotypes along our predicted pathway demonstrated an advantage in replication, fitness, or fidelity that would favor their maintenance or expansion in the viral population. Thus, natural selection is unlikely to drive ExoN-AA down these pathways towards reversion.

**Figure 2.**
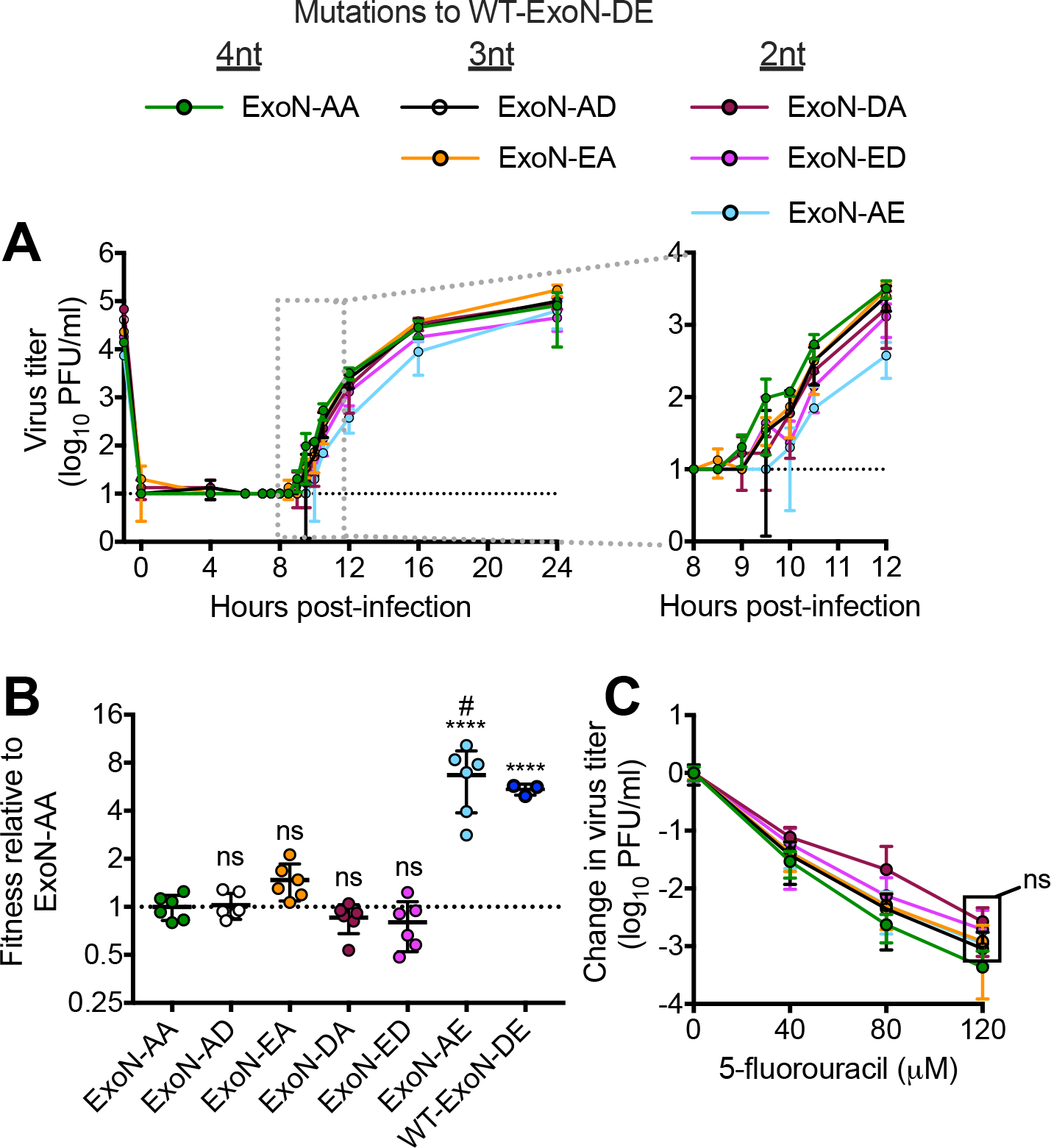
Intermediate revertants of ExoN-AA motif I do not have selective advantages. (*A*) Replication kinetics at MOI = 0.01 PFU/cell plotted as mean ± SD of n = 3. (*B*) Competitive fitness of each variant relative to ExoN-AA. Viruses were competed with a tagged ExoN-AA-reference strain, and relative fitness was normalized to the mean of ExoN-AA. (*C*) 5-fluorouracil sensitivity at MOI = 0.01 PFU/cell. Statistical significance of each variant relative to ExoN-AA was determined by one-way ANOVA with multiple comparisons (Panel *D*) two-way ANOVA with Dunnett’s multiple comparisons (panel *C*). ****p<0.0001; ns = not significant. Data in (*B*) and (*C*) represent mean ± SD of n = 6. Boxed values have the same significance. ^#^All lineages of ExoN-AE reverted to WT-ExoN-DE during the experiment.

### Secondary adaptations outside of ExoN-AA motif I increase fitness along alternative pathways

Although we did not find fitness advantages to intermediate revertants, we also did not identify profound fitness costs that would drive their immediate loss from populations. We have previously demonstrated that during 250 passages (P250), ExoN-AA can adapt for increased replication, fitness, and fidelity via secondary mutations outside of motif I (7). We tested whether secondary adaptive mutations could exceed the fitness of ExoN-AA and its intermediate revertants. To examine the early adaptation of ExoN-AA, we studied passage 10 from the P250 passage series (Figure 3). ExoN-AA-P10 retains the ExoN-AA motif I genotype but has increased replication and reduced susceptibility to 5-FU, altogether manifesting in greater relative fitness (Figure 3) (7). We identified only six total mutations within ExoN-AA-P10 by dideoxy sequencing (Table 2), indicating that rapid adaptation of and compensation for ExoN-AA requires relatively few genetic changes at the consensus level. To test whether interactions between multiple mutations or population level effects contribute to ExoN-AA-P10 fitness, we isolated a plaque-purified clone of ExoN-AA-P10. The clone replicated to higher titers than the ExoN-AA-P10 population but had identical 5-FU sensitivity and relative fitness (Figure 3), indicating that genomes derived from a single virus plaque encode the adaptive changes required by the total population. Together, these data demonstrate that mutations outside of ExoN(-) motif I can confer greater fitness advantages than intermediate revertants even at early passages. These early adaptive mutations likely reduce the selective pressure for motif I reversion and place the intermediate revertants at a selective disadvantage.

**Table 2.** Mutations in ExoN(-) P10. Data derived from dideoxy sequencing.

<sup>a</sup>Mutation present at approximately 50% of population. <sup>b</sup>MHV HE is not transcribed in tissue culture. <sup>c</sup>Amino acid numbers designate positions within cleaved nsps, not the polyprotein.
| Mutation number | Nucleotide change | Protein | Amino acid change<sup>c</sup> |
| --- | --- | --- | --- |
| 1 | G2520A<sup>a</sup> | nsp2 | D524N |
| 2 | A3080G<sup>a</sup> | nsp3 | silent |
| 3 | T16017A | nsp12 | M814K |
| 4 | A17836G<sup>a</sup> | nsp13 | I492M |
| 5 | G22673A<sup>a</sup> | HE<sup>b</sup> | noncoding |
| 6 | A29298C<sup>a</sup> | M | silent |

**Figure 3.**
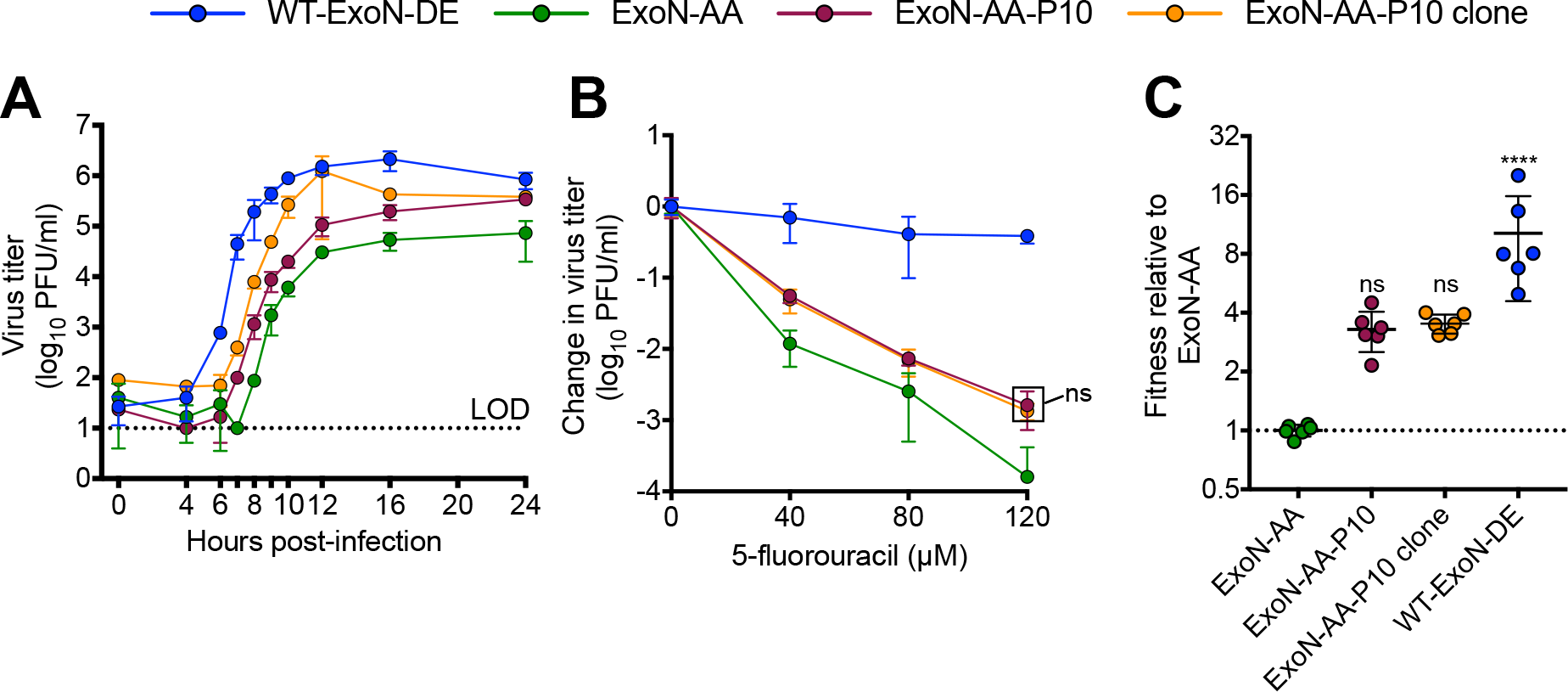
ExoN-AA adapts for increased fitness within 10 passages. (*A*) Replication kinetics of indicated viruses at MOI = 0.01 PFU/cell plotted as mean ± SD of n = 3. (*B*) 5-fluorouracil sensitivity at MOI = 0.01 PFU/cell. (*C*) Competitive fitness of individual recombinants relative to ExoN-AA. Viruses were competed with a tagged ExoN-AA-reference strain, and relative fitness was normalized to the mean of ExoN-AA. Statistical significance of each virus relative to ExoN-AA was determined by two-way ANOVA with Dunnett’s multiple comparisons (panel *B*) or by one-way ANOVA with multiple comparisons (Panel *C*). ****p < 0.0001, ns = not significant. LOD = limit of detection. Data in (*B*) and (*C*) represent mean ± SD of n = 6. Boxed values have the same significance.

### Adaptive mutations in nsp12 and nsp14 that increase ExoN-AA fitness confer significant fitness costs to WT-ExoN-DE

Mutational fitness effects are highly dependent upon the genetic background (25–27). In addition to reducing selective pressure for reversion, mutations conferring increased fitness to ExoN-AA might also reduce the benefits of motif I reversion. We previously reported that long-term passage of ExoN-AA selects for secondary adaptive mutations in the nsp12 RdRp and nsp14 (nsp12-P250 and nsp14-P250) (7). Nsp12-P250 contains 7 nonsynonymous mutations that partially compensate for defective proofreading and increase ExoN-AA fitness. Nsp14-P250 contains 6 nonsynonymous mutations, including a conservative D-to-E substitution in ExoN motif III, and increases ExoN-AA fitness without compensating for defective proofreading. To test whether the fitness effects of passage-associated mutations in nsp12-P250 and nsp14-P250 depend upon the ExoN-AA genotype, we engineered a WT motif I (ExoN-DE) into viruses containing nsp12-P250 and nsp14-P250, alone and together, and analyzed replication, 5-FU sensitivity, and competitive fitness. Compared to WT-ExoN-DE, both ExoN-DE-nsp12-P250 and ExoN-DE-nsp14-P250 displayed delayed and decreased replication (Figure 4A). In 5-FU sensitivity assays, ExoN-DE-nsp14-P250 was indistinguishable from WT-ExoN-DE, while both variants containing nsp12-P250 (ExoN-DE-nsp12-P250 and ExoN-DE-nsp12/14-P250) were significantly more sensitive to 5-FU (Figure 4B). Finally, the nsp12-P250 and nsp14-P250 mutations significantly decreased fitness relative to WT-ExoN-DE (Figure 4C). We detected no statistical differences between the specific infectivity of WT-ExoN-DE and any of the nsp12-P250 and nsp14-P250 variants in isolated infections (Figure 4D). Thus, mutations in nsp12 and nsp14 that arose in the ExoN-AA background were detrimental to replication, mutagen sensitivity, and competitive fitness in the presence of a fully-reverted ExoN-DE. These results support the conclusion that the adaptive pathways available to ExoN-AA may stabilize the ExoN-AA genotype, reducing both the selective pressure for, and the potential benefits of, primary reversion.

**Figure 4.**
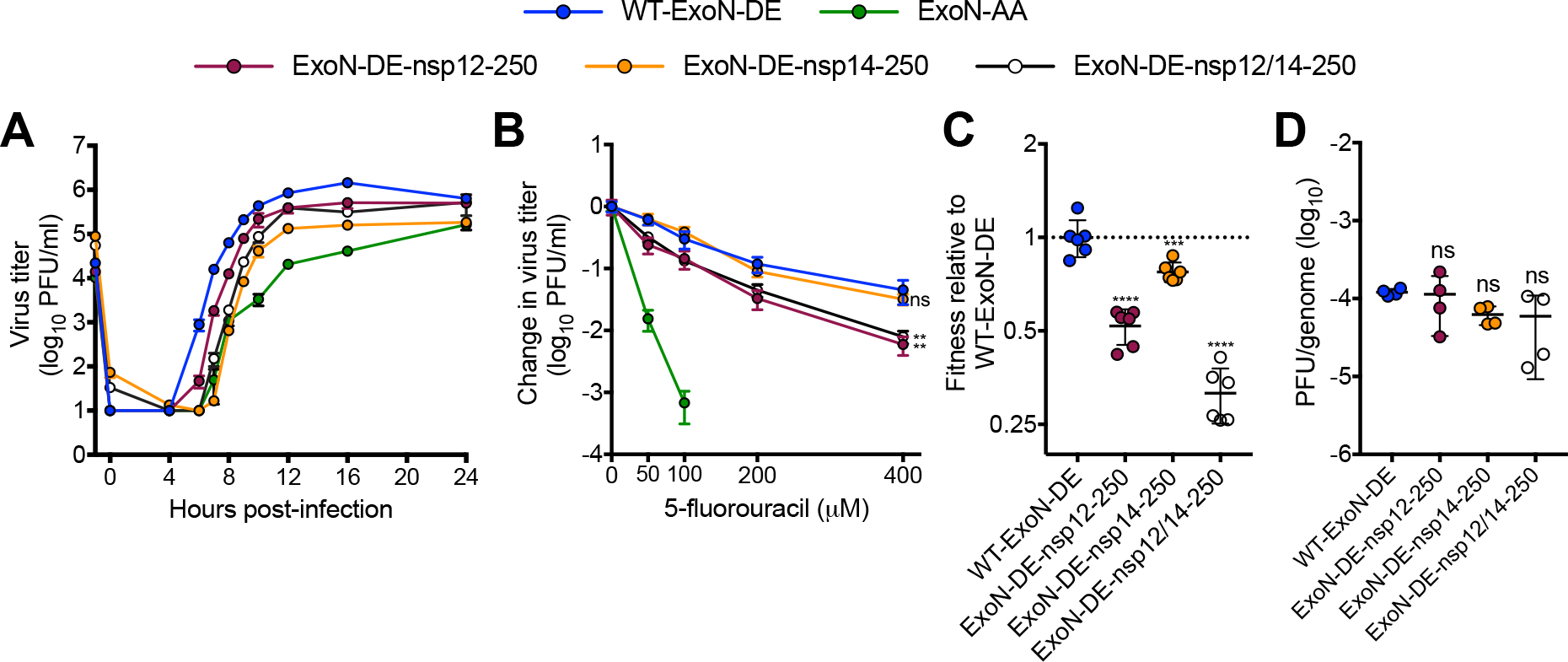
Mutations that increase ExoN-AA fitness are detrimental in the presence of WT-ExoN-DE. (*A*) Replication kinetics of indicated viruses at MOI = 0.01 PFU/cell plotted as mean ± SD of n = 3. (*B*) 5-fluorouracil sensitivity at MOI = 0.01 PFU/cell, mean ± SD of n = 6. (*C*) Competitive fitness of individual recombinants relative to WT-ExoN-DE. Viruses were competed with a tagged WT-ExoN-DE reference strain, and relative fitness was normalized to the mean of WT-ExoN-DE, mean ± SD of n = 6. (*D*) Specific infectivity (genomes per PFU) from isolated infections, mean ± SD of n = 4.. Statistical significance of each virus relative to WT-ExoN-DE was determined with two-way ANOVA with Dunnett’s multiple comparisons test (panel *B*) or by ordinary one-way ANOVA with Dunnett’s multiple comparisons test (panels *C* and *D*). **p < 0.01; ***p < 0.001; ****p < 0.0001; ns = not significant.

## DISCUSSION

In this study, we demonstrate that the stability of the ExoN(-) motif I genotype in MHV (ExoN-AA) is a consequence of the limitations and opportunities of the genetic landscape it explores during replication (Figure 5). Our results support a model in which the viable adaptive pathways leading to direct reversion of motif I from AA-to-DE are relatively flat on a fitness landscapes, as intermediate revertants remain phenotypically ExoN(-) and confer no fitness advantage over ExoN-AA. In contrast, at least one alternative adaptive pathway is readily accessible and imparts immediate fitness gains over ExoN-AA. We propose that even minimal alternative pathway adaptive fitness gains reduce the likelihood and benefits of motif I reversion, until eventually the changing genetic background renders reversion detrimental. These data and this model suggest that selection during replication favors immediate, incremental fitness gains along the most accessible pathway rather than dramatic fitness increases across a larger genetic barrier.

**Figure 5.**
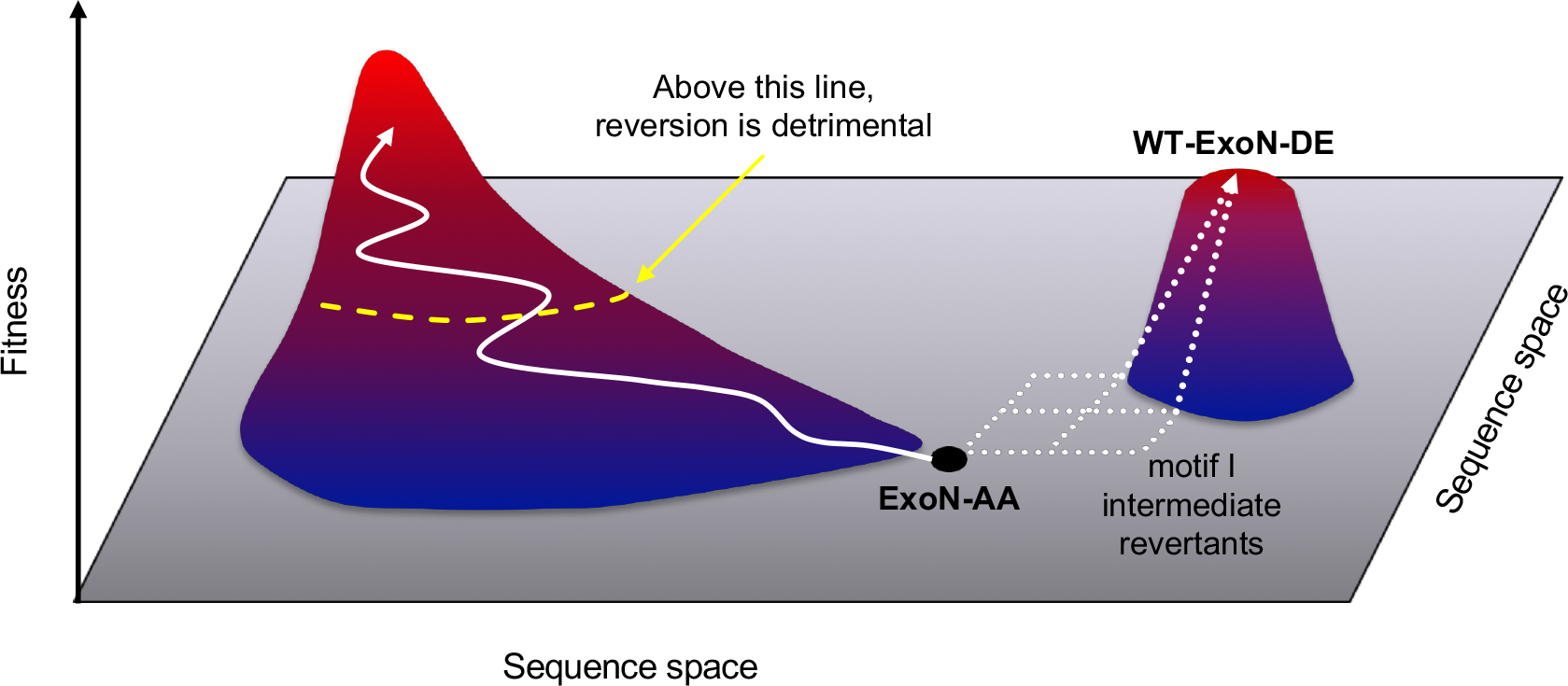
Model for the *in vitro* evolution of MHV-ExoN-AA. MHV-ExoN-AA(black dot) is a low-fitness variant. Reversion to WT-ExoN-DE would dramatically increase fitness but can only be achieved by traversing a flat landscape and climbing a steep fitness cliff (dotted white arrows). However, secondary mutations that incrementally increase fitness are more accessible (solid white arrow). Eventually, the genetic background changes enough that reversion becomes detrimental (dotted yellow line).

Our results also extend existing studies of CoV ExoN motif I. Motif I AA→DE mutations in the SARS-CoV nsp14-ExoN dramatically reduce nuclease activity in biochemical assays, but no study has examined the contributions of each residue independently (16, 18). Intermediate revertants of ExoN-AA did not display consistent or statistical differences in replication, 5-FU sensitivity, or competitive fitness relative to ExoN-AA, suggesting that they remain phenotypically ExoN(-) during infection and supporting previous studies that motif I DE is essential for WT ExoN function. Given these results, we were surprised to observe repeated reversion of the ExoN-AE but not the other two 2nt variants, ExoN-DA and ExoN-ED. One potential explanation is that the specific mutational bias of ExoN-AE makes the revertant mutations more accessible than in ExoN-DA or ExoN-ED. Alternatively, if ExoN-AE has profound replication or fitness defects, selection could drive primary reversion more quickly away from this genotype. Consistent with this hypothesis, ExoN-AE reverted more quickly at a higher MOI, where natural selection acts more efficiently on a larger population size (Table 1) (28). Biochemical studies would be valuable to understand how nsp14-ExoN-AE differs from the other intermediate revertants, with an eye towards understanding the catalytic constraints and functional interactions with nascent CoV RNA.

CoV replication proceeds through the concerted action of multiple proteins proposed to resemble DNA replication holoenzymes (29). Due to the extensive interactions between CoV proteins, they must coevolve in a highly cooperative manner to maintain their essential functions. Consistent with this hypothesis, the fitness effects of mutations in nsp12-P250 and nsp14-P250 differ based on the motif I genotype; they are beneficial in ExoN-AA but detrimental in the WT-ExoN-DE background. In previous studies, it has been difficult to determine whether the fitness defects in ExoN(-) CoVs are directly linked to low-fidelity replication or through some other mechanism. Our data suggest that the proofreading function of nsp14-ExoN can be uncoupled from its more general role in replication (Figure 4), providing an opportunity to examine additional roles for this essential protein. Nsp12-P250 also will be an important tool for understanding the relationship between RdRp fidelity and ExoN proofreading during CoV replication and for studying replication complex assembly and interactions. Our studies suggest that compensatory mutations identified through long-term passage could stabilize the ExoN-AA genotype. In particular, the high-fidelity nsp12-P250 could reduce the probability of reversion by reducing mutational sampling within motif I (30), and both nsp12-P250 and nsp14-P250 render the MHV genome inhospitable to a WT-ExoN-DE. Together, these studies argue that experimental evolution can generate reagents to define critical interactions involved in CoV replication and can identify new strategies for stabilizing attenuated CoVs.

## MATERIALS AND METHODS

### Cell culture

Delayed brain tumor (DBT-9) cells (31) and baby hamster kidney 21 cells expressing the MHV receptor (BHK-R) (32) were maintained at 37°C in Dulbecco’s Modified Eagle Medium (DMEM, Gibco) supplemented with 10% serum (HyClone FetalClone II, GE Healthcare or Fetal Bovine Serum, Invitrogen), 100 U/mL penicillin and streptomycin (Gibco), and 0.25 μM amphotericin B (Corning). BHK-R cells were further supplemented with 0.8 mg/mL G418 selection antibiotic (Gibco). The infectious clone of the murine hepatitis virus strain A59 (MHV-A59; GenBank accession number AY910861) was used as the template for all recombinant viruses.

### Determination of viral titer by plaque assay

Virus samples were serially diluted and inoculated on subconfluent DBT-9 cell monolayers in either 6- or 12-well format. Cells were overlaid with 1% agar in DMEM and incubated overnight at 37°C. Plates were fixed with 4% formaldehyde and agar plugs were removed. The number of plaques per well was counted by hand and used to calculate titer (32).

### Plaque purification of viral populations

DBT cells were infected with serial dilutions of virus and overlaid with 1% agar in DMEM. Single plaques were isolated with glass Pasteur pipettes, resuspended in PBS containing calcium and magnesium, and inoculated onto fresh DBTs. This process was completed 3 times before generating experimental stocks.

### Cloning and recovery of recombinant viruses

Site-directed mutagenesis in MHV genome fragments was performed using “round the horn” PCR (originally described in (33)). Briefly, adjacent primers containing the mutation of interest were 5′-phosphorylated using T4 polynucleotide kinase (NEB, M0201S) using the buffer from the T4 DNA ligase, which contains ATP (M0202S). PCR was performed on a plasmid template using the Q5 High-fidelity 2× Master Mix (NEB, M0492L), with primers at final concentration of 500nM. The linear amplification product was purified using the Promega Wizard SV Gel and PCR Clean-up System (Promega Corporation, A9282), and 4 μL was ligated at 16°C overnight with the T4 DNA ligase (NEB M0202S). After transformation into chemically-competent Top10 *E. coli* (lab-derived) and expansion in liquid culture, the MHV segment of each plasmid was sequenced. Viruses were constructed, rescued, and sequenced as described previously (7, 13, 32). Experimental stocks were generated by infecting a subconfluent 150 cm^2^ flask of DBT-9 cells at MOI of 0.01 PFU/cell. Flasks were frozen at −80°C when monolayers were fully involved, approximately 20-28 hours post-infection depending on the variant. After thawing, the supernatant was clarified by centrifugation at 4,000 x g (Sorvall RC 3B Plus; HA-6000A rotor) for 10 min at 4°C. For intermediate revertants, stocks were generated in serum-free DMEM and processed as above before being concentrated roughly 10-fold by centrifugation at 4,000 x g using Amicon Ultra-15 Centrifugal Filter Units, 100kDa (EMD Millipore, UFC910008). The virus titer of each stock was determined by plaque assay using DBT-9 cells as described above.

### Passage of ExoN intermediate revertants

Intermediate revertants of ExoN-AA were passaged 10 times on subconfluent DBT-9 cell monolayers in 24-well plates at an estimated MOI of either 0.01 or 0.5 PFU/cell. Supernatants were harvested at 24 and 20 hours post-infection for MOI = 0.01 and 0.5 PFU/cell, respectively, and screened for WT reversion by plaque assay. At least three WT-like plaques were sequenced for each lineage to confirm motif I reversion.

### Replication kinetics

Viral replication kinetics in DBT-9 cells were determined at indicated MOIs as described previously (11). Replicates were synchronized by 30-minute incubation at 4°C before transferring to the 37°C incubator. Supernatant (300 μL) was harvested at the indicated time points and titered by plaque assay.

### Determination of specific infectivity

Subconfluent monolayers of DBT-9 cells in 24-well plates were infected with the indicated virus at MOI = 0.05 PFU/cell, and supernatant was harvested at 16 hours post-infection. Genomic RNA in supernatant was quantified using one-step reverse transcription quantitative RT-PCR (RT-qPCR) on TRIzol-extracted RNA as described previously (9). Briefly, genomic RNA was detected with a 5’ 6-carboxyfluorescein (FAM) and 3’ black hole quencher 1 (BHQ-1) labeled probe targeting nsp2 (Biosearch Technologies, Petaluma, CA), and RNA copy number was calculated by reference to an RNA standard derived from the MHV A fragment. Samples were plated in technical duplicate to minimize well-to-well variation. Titers were determined by plaque assay in DBT-9 cells, and specific infectivity was calculated as PFU per supernatant genomic RNA copy.

### 5-fluorouracil sensitivity assays

Stock solutions of 5-fluorouracil (Sigma F6627) were prepared in dimethyl sulfoxide (DMSO). Sensitivity assays were performed in 24-well plates at MOI = 0.01 PFU/cell, as previously described (7). Cells were incubated with drug for 30 minutes prior to infection. Supernatants were harvested at 24 hours post-infection, and titers were determined by plaque assay.

### Competitive fitness assays

ExoN-AA-reference and WT-ExoN-DE-reference viruses were marked with 7 consecutive silent mutations within nsp2 (wild-type: 1301-TTCGTCC-1307; reference: 1301-CAGCAGC-1307) by round the horn PCR, as described above. Competitions were performed in triplicate on DBT-9 cells in 12-well plates, plated at a density of 1 × 10^5^ cells per well 24 hours prior to infection. Cells were infected at a total MOI of 0.1 PFU/cell (MOI = 0.05 PFU/cell each for competitor and reference virus). Supernatants were harvested 15 and 16 hours post-infection for experiments with ExoN-AA-reference and WT-ExoN-DE-reference, respectively, and passaged 4 times. Samples were titered between all passages to maintain total MOI of 0.1 PFU/cell. RNA was extracted from 70 μL of supernatant using QIAamp 96 virus QIAcube HT kit on the QIAcube HT System (Qiagen). Each RNA sample was analyzed by one-step RT-qPCR with two SYBR Green assays. Reference viruses were detected with forward primer SS-qPCR-Sil-F (5′-CTATGCTGTATACGGA<u>CAGCAGT</u>-3′; 200nM final) and reverse primer SS-qPCR-R2 (5′-GGTGTCACCACAACAATCCAC-3′, 200nM final). Competitors were detected with forward primer SS-qPCR-WT-F (5′-CTATGCT-GTATACGGA<u>TTCGTCC</u>-3′, 450 nM final) and reverse primer SS-qPCR-R2 (5′-GGTGTCAC-CACAACAATCCAC-3′, 450 nM final). RNA samples were diluted 1:100 prior to RT-qPCR with *Power* SYBR Green RNA-to-Ct 1-step kit (Applied Biosystems) according to the manufacturer’s protocol. Duplicate wells were averaged, and values were excluded from subsequent analysis if the duplicate wells differed by > 0.5 Ct. The relative abundance of competitor and reference were determined by subtracting Ct thresholds (ΔCt_competitor_ = Ct_competitor_ – Ct_reference_) and converted to reflect the fold-change in ratio (Δratio = 2^−ΔCt competitor^). The log10Δratio was plotted against passage number, and the change in log_10_Δratio (i.e. slope of linear regression) is the relative fitness. Note that regressions were fit only through P1-P4, as slight deviations in 1:1 ratio in the input (P0) can skew the slope.

### Statistical analysis

GraphPad Prism 6 (La Jolla, CA) was used to perform statistical tests. Only the comparisons shown [e.g. ns or asterisk(s)] within the figure or legend were performed. In many cases the data were normalized to untreated controls. This was performed using GraphPad Prism 6. The number of replicate samples is denoted within each figure legend.

## ACKNOWLEDGEMENTS

We thank members of the Denison laboratory and Seth Bordenstein for valuable discussions, as well as Andrea Pruijssers for critical review of the manuscript. This work was supported by United States Public Health Service awards R01-AI108197 (M.R.D), T32-GM007347 (K.W.G), F30-AI129229 (K.W.G), T32-AI089554 (N.R.S.), F31-AI133952 (M.L.A.), and T32-AI089554 (M.L.A.) all from the National Institutes of Health. The content is solely the responsibility of the authors and does not necessarily represent the official views of the National Institutes of Health.

The authors declare no conflicts of interest.

## REFERENCES

1. Sanjuán R, Nebot MR, Chirico N, Mansky LM, Belshaw R. 2010. Viral mutation rates. J Virol 84:9733–9748.

2. Domingo E, Sheldon J, Perales C. 2012. Viral quasispecies evolution. Microbiol Mol Biol Rev 76:159–216.

3. Dolan PT, Whitfield ZJ, Andino R. 2018. Mapping the Evolutionary Potential of RNA Viruses. Cell Host and Microbe 23:435–446.

4. Stern A, Yeh Te M, Zinger T, Smith M, Wright C, Ling G, Nielsen R, Macadam A, Andino R. 2017. The Evolutionary Pathway to Virulence of an RNA Virus. Cell 169:35–35.e19.

5. Perlman S, Netland J. 2009. Coronaviruses post-SARS: update on replication and pathogenesis. Nat Rev Microbiol 7:439–450.

6. Agostini ML, Andres EL, Sims AC, Graham RL, Sheahan TP, Lu X, Smith EC, Case JB, Feng JY, Jordan R, Ray AS, Cihlar T, Siegel D, Mackman RL, Clarke MO, Baric RS, Denison MR. 2018. Coronavirus Susceptibility to the Antiviral Remdesivir (GS-5734) Is Mediated by the Viral Polymerase and the Proofreading Exoribonuclease. MBio 9:e00221–18–15.

7. Graepel KW, Lu X, Case JB, Sexton NR, Smith EC, Denison MR. 2017. Proofreading-Deficient Coronaviruses Adapt for Increased Fitness over Long-Term Passage without Reversion of Exoribonuclease-Inactivating Mutations. MBio 8:e01503–17.

8. Smith EC, Blanc H, Surdel MC, Vignuzzi M, Denison MR. 2013. Coronaviruses lacking exoribonuclease activity are susceptible to lethal mutagenesis: evidence for proofreading and potential therapeutics. PLoS Pathog 9:e1003565.

9. Sexton NR, Smith EC, Blanc H, Vignuzzi M, Peersen OB, Denison MR. 2016. Homology-Based Identification of a Mutation in the Coronavirus RNA-Dependent RNA Polymerase That Confers Resistance to Multiple Mutagens. J Virol 90:7415–7428.

10. Case JB, Li Y, Elliott R, Lu X, Graepel KW, Sexton NR, Smith EC, Weiss SR, Denison MR. 2017. Murine Hepatitis Virus nsp14 Exoribonuclease Activity Is Required for Resistance to Innate Immunity. J Virol 92:e01531–17–38.

11. Smith EC, Case JB, Blanc H, Isakov O, Shomron N, Vignuzzi M, Denison MR. 2015. Mutations in coronavirus nonstructural protein 10 decrease virus replication fidelity. J Virol 89:6418–6426.

12. Graham RL, Becker MM, Eckerle LD, Bolles M, Denison MR, Baric RS. 2012. A live, impaired-fidelity coronavirus vaccine protects in an aged, immunocompromised mouse model of lethal disease. Nat Med 18:1820–1826.

13. Eckerle LD, Lu X, Sperry SM, Choi L, Denison MR. 2007. High fidelity of murine hepatitis virus replication is decreased in nsp14 exoribonuclease mutants. J Virol 81:12135–12144.

14. Eckerle LD, Becker MM, Halpin RA, Li K, Venter E, Lu X, Scherbakova S, Graham RL, Baric RS, Stockwell TB, Spiro DJ, Denison MR. 2010. Infidelity of SARS-CoV Nsp14-exonuclease mutant virus replication is revealed by complete genome sequencing. PLoS Pathog 6:e1000896.

15. Menachery VD, Gralinski LE, Mitchell HD, Dinnon KH III, Leist SR, Yount BL Jr., McAnarney ET, Graham RL, Waters KM, Baric RS. 2018. Combination attenuation offers strategy for live-attenuated coronavirus vaccines. J Virol JVI.00710–18–35.

16. Ma Y, Wu L, Shaw N, Gao Y, Wang J, Sun Y, Lou Z, Yan L, Zhang R, Rao Z. 2015. Structural basis and functional analysis of the SARS coronavirus nsp14-nsp10 complex. Proc Natl Acad Sci USA 112:9436–9441.

17. Snijder EJ, Bredenbeek PJ, Dobbe JC, Thiel V, Ziebuhr J, Poon LLM, Guan Y, Rozanov M, Spaan WJM, Gorbalenya AE. 2003. Unique and conserved features of genome and proteome of SARS-coronavirus, an early split-off from the coronavirus group 2 lineage. J Mol Biol 331:991–1004.

18. Bouvet M, Imbert I, Subissi L, Gluais L, Canard B, Decroly E. 2012. RNA 3’-end mismatch excision by the severe acute respiratory syndrome coronavirus nonstructural protein nsp10/nsp14 exoribonuclease complex. Proc Natl Acad Sci USA 109:9372–9377.

19. Minskaia E, Hertzig T, Gorbalenya AE, Campanacci V, Cambillau C, Canard B, Ziebuhr J. 2006. Discovery of an RNA virus 3“->5” exoribonuclease that is critically involved in coronavirus RNA synthesis. Proc Natl Acad Sci USA 103:5108–5113.

20. Becares M, Pascual-Iglesias A, Nogales A, Sola I, Enjuanes L, Zuñiga S. 2016. Mutagenesis of Coronavirus nsp14 Reveals Its Potential Role in Modulation of the Innate Immune Response. J Virol 90:5399–5414.

21. Steitz TA, Steitz JA. 1993. A general two-metal-ion mechanism for catalytic RNA. Proc Natl Acad Sci USA 90:6498–6502.

22. Chen P, Jiang M, Hu T, Liu Q, Chen XS, Guo D. 2007. Biochemical characterization of exoribonuclease encoded by SARS coronavirus. J Biochem Mol Biol 40:649–655.

23. Derbyshire V, Grindley ND, Joyce CM. 1991. The 3“-5” exonuclease of DNA polymerase I of Escherichia coli: contribution of each amino acid at the active site to the reaction. EMBO J 10:17–24.

24. Case JB, Ashbrook AW, Dermody TS, Denison MR. 2016. Mutagenesis of S-Adenosyl-l-Methionine-Binding Residues in Coronavirus nsp14 N7-Methyltransferase Demonstrates Differing Requirements for Genome Translation and Resistance to Innate Immunity. J Virol 90:7248–7256.

25. Das SR, Hensley SE, Ince WL, Brooke CB, Subba A, Delboy MG, Russ G, Gibbs JS, Bennink JR, Yewdell JW. 2013. Defining Influenza A Virus Hemagglutinin Antigenic Drift by Sequential Monoclonal Antibody Selection. Cell Host and Microbe 13:314–323.

26. Nakajima K, Nobusawa E, Nagy A, Nakajima S. 2005. Accumulation of Amino Acid Substitutions Promotes Irreversible Structural Changes in the Hemagglutinin of Human Influenza AH3 Virus during Evolution. J Virol 79:6472–6477.

27. Koel BF, Burke DF, van der Vliet S, Bestebroer TM, Rimmelzwaan GF, Osterhaus ADME, Smith DJ, Fouchier RAM. 2018. Epistatic interactions can moderate the antigenic effect of substitutions in hemagglutinin of influenza H3N2 virus 1–16.

28. Dolan PT, Whitfield ZJ, Andino R. 2018. Mechanisms and Concepts in RNA Virus Population Dynamics and Evolution. Annu Rev Virol 5:annurev–virology–101416–041718–24.

29. Smith EC, Sexton NR, Denison MR. 2014. Thinking Outside the Triangle: Replication Fidelity of the Largest RNA Viruses. Annu Rev Virol 1:111–132.

30. Arnold JJ, Vignuzzi M, Stone JK, Andino R, Cameron CE. 2005. Remote site control of an active site fidelity checkpoint in a viral RNA-dependent RNA polymerase. J Biol Chem 280:25706–25716.

31. Chen W, Baric RS. 1996. Molecular anatomy of mouse hepatitis virus persistence: coevolution of increased host cell resistance and virus virulence. J Virol 70:3947–3960.

32. Yount B, Denison MR, Weiss SR, Baric RS. 2002. Systematic assembly of a full-length infectious cDNA of mouse hepatitis virus strain A59. J Virol 76:11065–11078.

33. Ho SN, Hunt HD, Horton RM, Pullen JK, Pease LR. 1989. Site-directed mutagenesis by overlap extension using the polymerase chain reaction. Gene 77:51–59.

